# Decitabine reverses innate immune gene suppression in rare melanomas

**DOI:** 10.1101/2025.08.24.671992

**Authors:** Morgan L. MacBeth, Stacey M. Bagby, Jessica S.W. Borgers, Jacqueline A. Turner, Kelsey C. Anderson, Phaedra A. Whitty, Sarah Hartman, Beteleham Yacob, Vito W. Rebecca, Eduardo Davila, Todd M. Pitts, Theresa M. Medina, Sapna P. Patel, Martin D. McCarter, William A. Robinson, Richard P. Tobin, Kasey L. Couts

## Abstract

Rare melanoma subtypes, including acral, mucosal, and uveal melanomas, exhibit limited responses to immune checkpoint inhibitors (ICIs), yet the molecular mechanisms of immune resistance remain poorly defined. Here, we performed transcriptomic profiling of patient-derived xenografts (PDXs) and publicly available tumor datasets to systematically compare intratumoral gene expression across cutaneous and rare melanoma subtypes. We identified a convergent downregulation of innate immune pathogen sensing (IIPS) and type I interferon signaling pathways in rare melanomas compared to cutaneous, with lower expression also observed in anti-PD-1 non-responder tumors. CIBERSORT deconvolution of immune populations revealed that lower IIPS gene-expressing tumors exhibited reduced CD8⁺ T cell and memory CD4⁺ T cell infiltration, and enrichment of M2 macrophages, consistent with a more immunosuppressive tumor microenvironment. *In vitro* screening of epigenetic and immunomodulatory compounds revealed that the DNA hypomethylating agent decitabine robustly induced IIPS and adaptive immune gene expression in rare melanoma cell lines. *In vivo* treatment of mucosal and uveal melanoma xenograft models with decitabine resulted in durable upregulation of IIPS and antigen presentation genes, and whole transcriptome analysis confirmed that IIPS gene re-expression was the dominant transcriptional consequence of decitabine treatment. These findings highlight silencing of IIPS genes as a recurrent immune evasion mechanism in rare melanomas and nominate decitabine as a potential immunomodulatory strategy for enhancing immune responsiveness.

## Introduction

Immune checkpoint inhibitors (ICIs) have transformed the treatment landscape for cutaneous melanoma, where durable responses and long-term survival are now achievable for a substantial proportion of patients.^1,2^ However, this success has not extended equally across all melanoma subtypes. Acral, mucosal, and uveal melanomas (collectively referred to as rare melanomas) account for a minority of melanoma diagnoses but carry a disproportionate share of morbidity and mortality.^3,4^ These tumors exhibit significantly lower response rates to ICIs, and many rare melanoma tumors progress despite receiving standard-of-care immunotherapy.^5–14^

While several factors have been proposed to explain the limited efficacy of ICI in rare melanomas, including lower tumor mutational burden (TMB), reduced PD-L1 expression, and diminished T cell infiltration, these features vary considerably across subtypes. While TMB is consistently lower in acral, mucosal, and uveal melanomas compared to cutaneous melanoma,^15–18^ PD-L1 expression shows a more heterogeneous pattern, with similar levels in cutaneous and acral melanomas (∼35%), higher expression in mucosal melanoma (∼44%), and markedly lower expression in uveal melanoma (∼10%).^19,20^ Likewise, tumor-infiltrating lymphocyte (TIL) density is generally reduced in acral and mucosal melanomas, while uveal melanoma metastases appear to have TIL infiltration levels comparable to cutaneous disease.^21–24^ These inconsistencies highlight the limitations of current biomarkers in explaining immune resistance in rare melanoma subtypes. Moreover, no studies to date have systematically compared the transcriptional programs of acral, mucosal, and uveal melanomas to those of cutaneous melanoma. Without a clear molecular framework for understanding subtype-specific immune evasion, the development of effective therapeutic strategies remains a major challenge.

To address this gap, we undertook a comprehensive transcriptomic comparison of cutaneous, acral, mucosal, and uveal melanomas using both patient-derived xenograft models and publicly available tumor datasets. We identified low innate immune pathogen sensing (IIPS) pathway gene expression as a shared molecular feature potentially underlying immune resistance in rare melanomas. We further demonstrate that decitabine restores and enhances IIPS and adaptive immune gene expression, laying the groundwork for future therapeutic strategies that expand immunotherapy benefit to patients with these historically underserved melanoma subtypes.

## Methods

### Patient-derived xenograft models

All tissues and clinical data were collected by the University of Colorado Skin Cancer Biorepository from patients with informed consent under COMIRB protocol #05-0309. Due to the limited availability and variable RNA quality of fresh surgical melanoma specimens, we routinely establish patient-derived xenograft (PDX) models by subcutaneously implanting fresh tumor fragments into female athymic nude mice (Athymic Nude-Foxn1nu, Inotiv) as previously described.^25^ Tumors were either harvested for tissue collection or passaged into a new generation of mice for further expansion. All animal procedures were approved by and conducted in accordance with the guidelines of the University of Colorado Anschutz Medical Campus Institutional Animal Care and Use Committee (IACUC).

### Whole transcriptome sequencing of PDX models and TCGA data analysis

RNA was extracted from early-generation (F1–F5) flash-frozen PDX tumors using the RNeasy Mini Kit (Qiagen), following tissue homogenization with the TissueLyser II (Qiagen) and on-column DNase digestion. RNA samples were submitted to the Genomics Core at National Jewish Health (Denver, CO) for whole-transcriptome sequencing (see Supplementary Methods for details). FASTQ files from our samples, as well as uveal melanoma FASTQ files downloaded from cBioPortal and TCGA pan-cancer cutaneous melanoma FASTQ files, were processed as described in the Supplementary Methods.

### Innate Immune Pathogen Sensing (IIPS) Gene Signature Construction

To generate an IIPS gene signature, we selected 15 genes based on prior literature describing canonical type I interferon response and innate immune pathogen sensing (IIPS) pathways. Genes were further prioritized based on consistent co-expression patterns observed in our transcriptomic dataset across melanoma subtypes. For each gene, normalized expression values were mean-centered across all samples, log_2_-transformed, and then averaged per sample to compute a composite IIPS score. This score served as a proxy for relative IIPS pathway activity and was used in subsequent correlative analyses.

### Immune cell deconvolution using CIBERSORT

To estimate the relative abundance of immune cell populations within PDX tumors, bulk RNA-seq data were analyzed using the CIBERSORT algorithm (https://cibersort.stanford.edu/), applying the LM22 signature matrix to profile 22 human immune cell types. Input expression data were normalized to transcripts per million (TPM) prior to analysis. The CIBERSORT pipeline was run using 100 permutations, and only samples with significant deconvolution results (*P* < 0.05) were included in downstream comparisons between high and low IIPS score groups.

### Cell lines and treatments

Melanoma cell lines and short-term cultures were either established in-house from PDX models (MB3572, MB3883, MB4199, MB2724, MB4486, MB4667, MB2141, MB4628) or purchased from ATCC (MP-41, 92-1). Cells were maintained in RPMI-1640 medium supplemented with 10% fetal bovine serum (FBS) and 1% penicillin–streptomycin at 37°C in a humidified incubator with 5% CO₂. For treatment studies, 2.5 x 10⁴ or 5.0 x 10⁴ cells were plated onto 6 cm dishes 24 hours prior to drug exposure. All cell lines were routinely tested for mycoplasma contamination and authenticated by short tandem repeat (STR) profiling.

### Mouse xenograft treatment studies

Melanoma cell lines maintained in RPMI-1640 medium were mixed 1:1 with Cultrex (R&D Systems) and injected subcutaneously (5 x 10⁶ cells per injection) into both flanks of ∼10-week-old female athymic nude mice. Once tumors reached ∼100–300 mm³, mice were randomized into two treatment groups (n = 10 per group) and treated with either vehicle or a combination of tetrahydrouridine and decitabine. Tetrahydrouridine (4 mg/kg) and decitabine (10 μg per mouse) were prepared in PBS and administered subcutaneously in a 100 μL volume once daily for three consecutive days following randomization. Tetrahydrouridine was given 30 minutes prior to decitabine to enhance its bioavailability. Mice were monitored for treatment-related side effects, and body weights were recorded twice weekly for the duration of the study.

### RNA sequencing of decitabine-treated cells and xenograft tumors

RNA was extracted from cell lines and xenograft tumors using the RNeasy Mini Kit (Qiagen) with on-column DNase digestion. Biological replicate RNA samples were submitted for sequencing and data analysis to GENEWIZ (Azenta Life Sciences, Inc.), as described in the

## Supplementary Methods

### Quantitative Real-Time PCR Quantitative Real-Time PCR

Total RNA was extracted from frozen tumors and cell lines using the RNeasy Plus Mini Kit (Qiagen) with on-column DNase digestion. Tumor samples were homogenized using the TissueLyser II (Qiagen) prior to RNA isolation. Complementary DNA (cDNA) was synthesized from 500 ng of input RNA using the Verso cDNA Synthesis Kit (Thermo Fisher Scientific). Quantitative real-time PCR was performed using PowerUp SYBR Green Master Mix (Thermo Fisher Scientific) on a QuantStudio 3 Real-Time PCR System (Applied Biosystems). All reactions were run in triplicate. Primer sequences are provided in the Supplementary Methods.

## Results

### Transcriptome analysis reveals low expression of innate immune pathogen sensing genes in rare melanoma subtypes

Given the limited evidence for genomic factors that explain poor immune responses in rare melanomas, we sought to characterize the intratumoral transcriptomic landscapes of acral, mucosal, and uveal melanoma compared to cutaneous melanoma. We performed whole-transcriptome RNA sequencing on 45 melanoma patient-derived xenograft (PDX) tumors, including 24 cutaneous, 8 acral, and 13 mucosal tumors. These PDX data were analyzed in parallel with The Cancer Genome Atlas (TCGA) tumor datasets for cutaneous and uveal melanoma using the same bioinformatic pipeline.^26^

Within each dataset, differential gene expression analysis was performed by comparing each rare melanoma subtype to cutaneous melanoma, using a log₂ fold-change threshold of ≥ 1.5 and an adjusted *P*-value of < 0.05 (Figure 1, Supplementary Data 1). Across all comparisons, rare melanoma subtypes consistently exhibited more downregulated than upregulated genes: acral (172 downregulated vs. 84 upregulated), mucosal (1,519 vs. 545), and uveal (7,468 vs. 5,489) (Figure 1A-B). Pathway analysis of the upregulated gene sets (Supplementary Data 2) revealed minimal enrichment: zero pathways for acral, one for mucosal, and twelve for uveal melanoma (Supplementary Figure 1A). Expected lineage-associated gene expression patterns were observed, including *CRKL* and *GAB2* in acral melanoma and *RASGRP3* in uveal melanoma (Supplementary Figure 1).^27–29^ Only a small number of genes were consistently upregulated across all rare melanoma subtypes (Figure 1B, Supplementary Data 1), with *JAG2* and *CECR2* among the most notable (Supplementary Figure 1B). A greater number of genes showed overlapping upregulation in at least two subtypes (Figure 1B, Supplementary Figures 1C-E).

**Figure 1.**
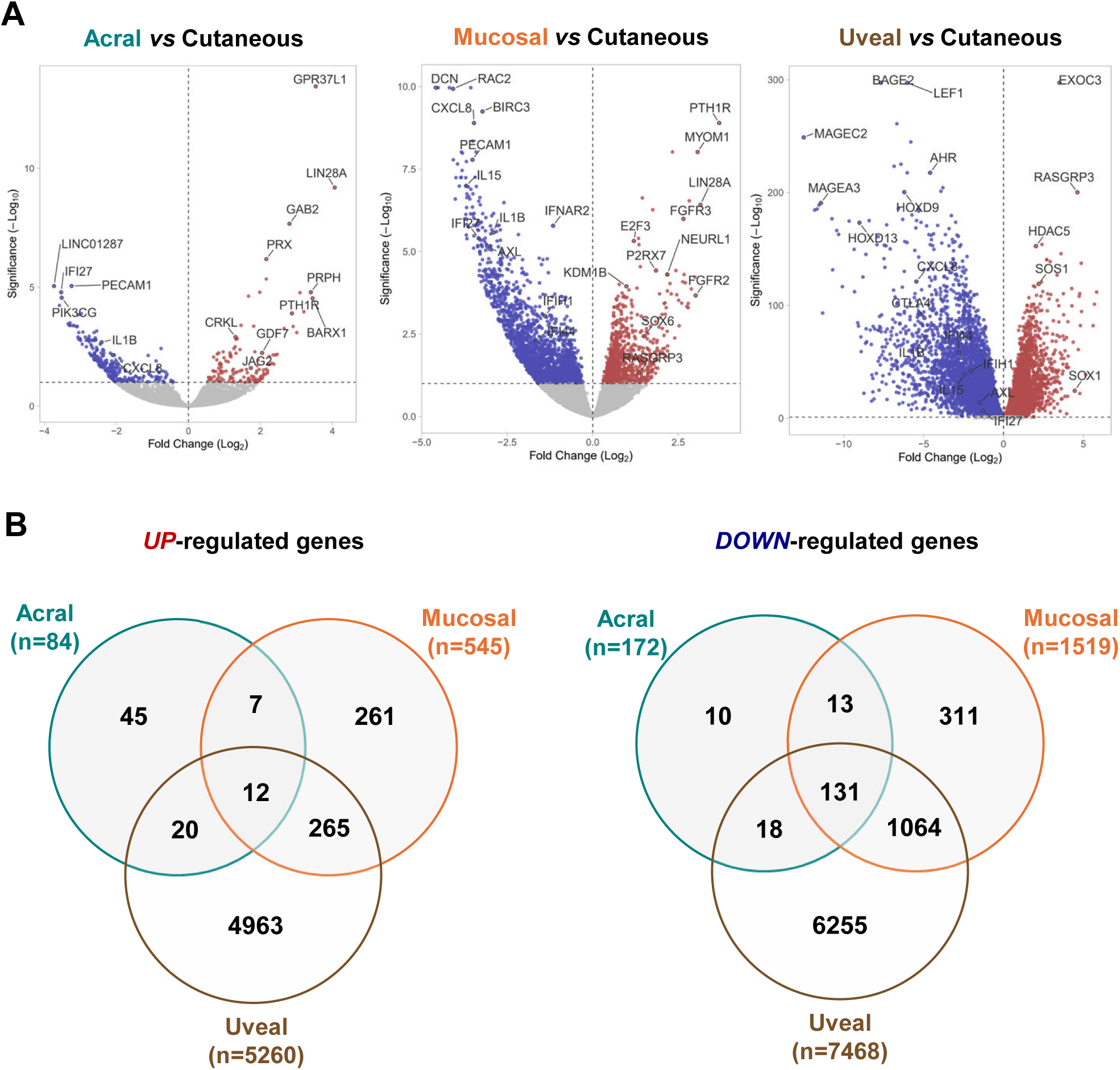
Differential gene expression analysis of rare melanomas compared to cutaneous melanoma. **(A)** Volcano plots showing differentially expressed genes for acral (left), mucosal (center), and uveal (right) melanomas relative to cutaneous melanoma. Genes significantly upregulated (Log₂ fold change ≥ 1.5; adjusted *P* < 0.05) are shown in red, and significantly downregulated genes in blue. Selected subtype-enriched or immune-related genes are labeled. **(B)** Venn diagrams illustrating the number of distinct and overlapping genes that are significantly upregulated (left) or downregulated (right) in each melanoma subtype compared to cutaneous melanoma.

In contrast to the limited number of upregulated genes, we identified numerous pathways enriched among the downregulated gene sets across rare melanoma subtypes (Figure 2A, Supplementary Data 2). Several of these pathways, and the genes within them (Supplementary Data 1), were consistently downregulated across all rare melanoma subtypes, including inflammatory response (e.g., *IL1B*, *CXCL8*), general immune signaling (e.g., *CTLA4*, *IL15*), and chemotaxis/migration pathways (e.g., *CCL20*, *CXCL10*) (Figure 2A-D). Other pathways showed more subtype-specific downregulation. For example, angiogenesis-related genes (*VEGFA*, *ANGPT2*) and genes involved in interferon (IFN) response and innate immune pathogen sensing (IIPS), including *IFNAR2*, *IFI44*, *IFI6*, and *CGAS*, were more preferentially suppressed in mucosal and uveal melanomas (Figure 2A, 2E-F).

**Figure 2.**
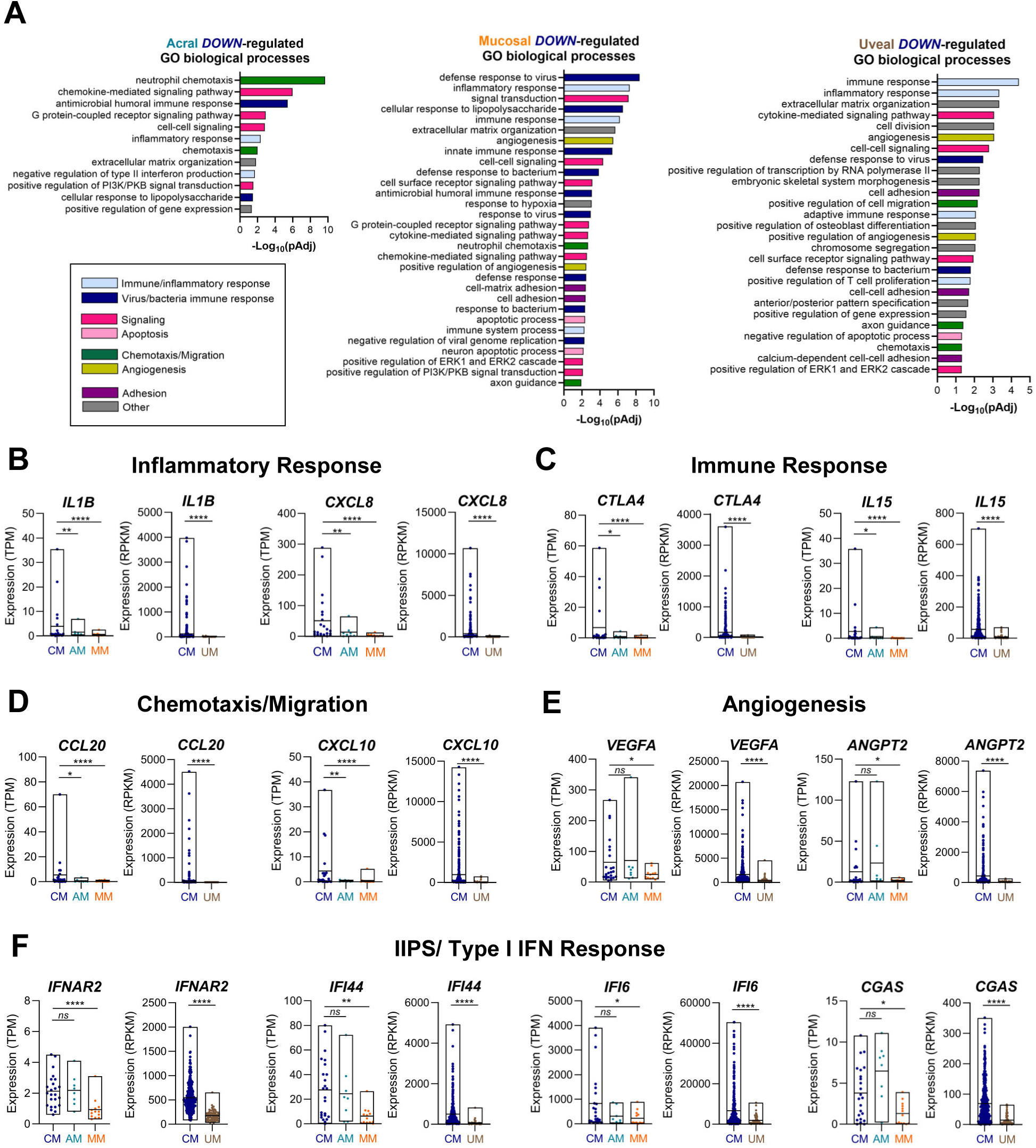
Downregulated immune and signaling pathways in rare melanomas. **(A)** Gene Ontology (GO) biological processes significantly enriched (*adjusted P* < 0.05) among genes downregulated in acral (left), mucosal (center), and uveal (right) melanomas compared to cutaneous melanoma. Pathways are color-coded by functional category (see legend). **(B–F)** RNA-seq expression of selected genes within downregulated pathways, including **(B)** inflammatory response (*IL1B*, *CXCL8),* **(C)** immune response (*CTLA4*, *IL15*), **(D)** chemotaxis/migration (*CCL20*, *CXCL10*), **(E)** angiogenesis (*VEGFA*, *ANGPT2*), and **(F)** innate immune pathogen-sensing/type I IFN signaling (*IFNAR2*, *IFI44*, *IFI6*, *CGAS*). PDX RNA-seq data are shown as TPM (transcripts per million); TCGA data are shown as FPKM (fragments per kilobase of transcript per million). Box plots display median and interquartile range. **** *P* < 0.0001, *** *P* < 0.001, ** *P* < 0.01, *P* < 0.05.

### Suppressed IIPS Gene Expression Correlates with Immune Exclusion and Anti-PD-1 Resistance

Among all enriched pathways, the IIPS pathway emerged as the most frequently and significantly downregulated across rare melanomas (Figure 2A). To evaluate its association with immunotherapy response, we curated a 15-gene IIPS signature and applied this to the PDX and TCGA RNA-seq datasets. Expression of IIPS genes was broadly reduced in rare melanoma subtypes compared to cutaneous melanoma (Figure 3A-B, Supplementary Figure 2A-C), and IIPS scores were lower in non-responders versus responders to anti-PD-1, or combination anti-CTLA-4 plus anti-PD-1, with significance being reached in rare melanoma patients (Figure 3C). Using CIBERSORT analysis, we further profiled immune cell populations within PDX tumors and found that tumors with lower IIPS scores (bottom 20%) were enriched for M2 macrophages, while those with higher IIPS scores (top 20%) exhibited increased CD8⁺ T cells, active memory CD4⁺ T cells, and M1 macrophages (Figure 3D). These findings were validated using TCGA RNA-seq data, where uveal melanomas showed consistently lower IIPS gene expression compared to cutaneous melanomas (Supplementary Figure 2A-C). CIBERSORT analysis of TCGA cutaneous melanoma samples further confirmed that tumors with higher IIPS scores (top 20%) harbored significantly greater CD8⁺ and memory CD4⁺ T cells, and fewer M2 macrophages (Supplementary Figure 2D). Collectively, these data suggest that IIPS pathway suppression is associated with unfavorable immune cell infiltration and resistance to ICI, supporting its potential as both a biomarker and therapeutic target in rare melanomas.

**Figure 3.**
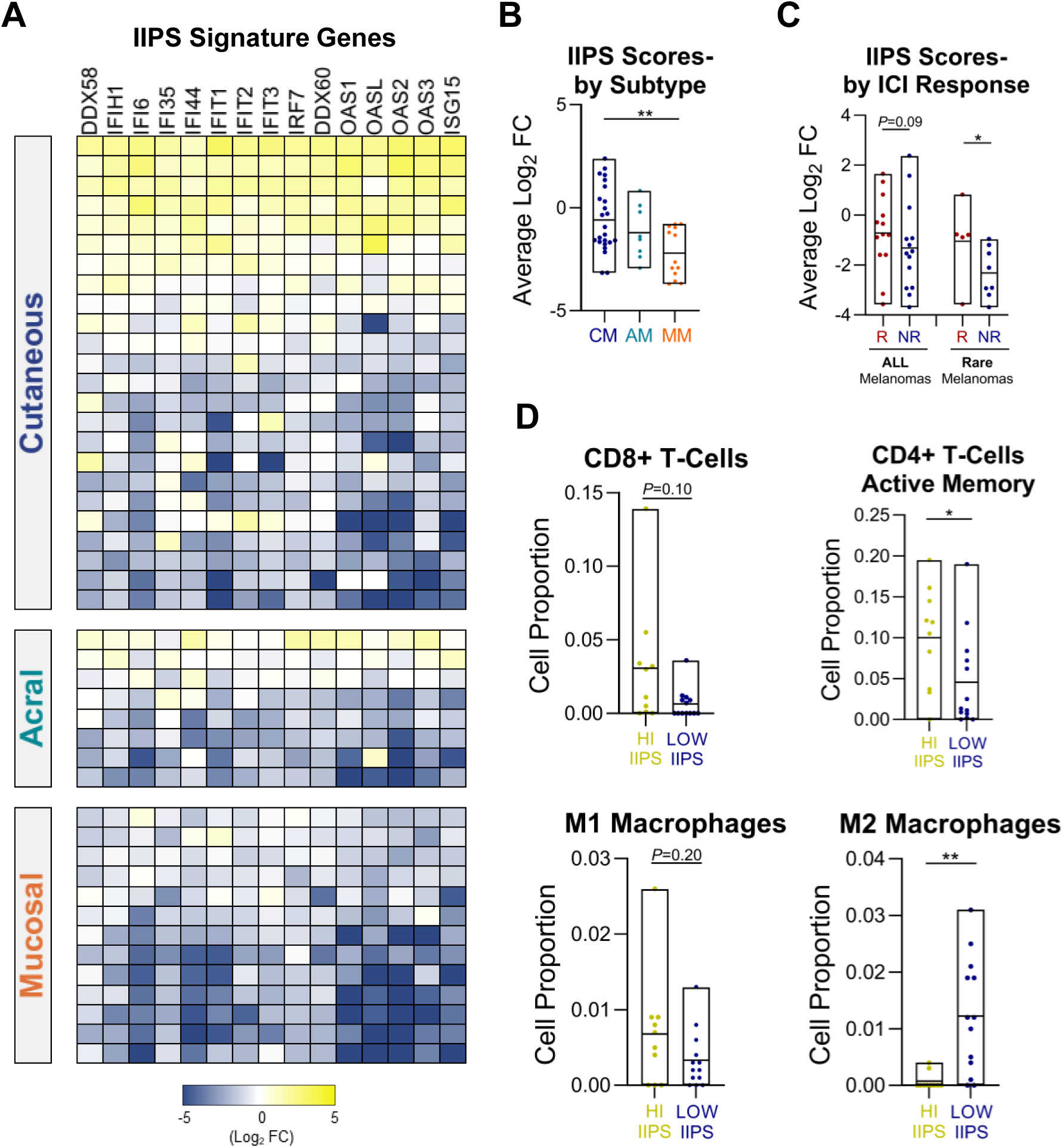
Low IIPS scores in rare melanoma PDX models are associated with reduced immune infiltration and worse response to anti-PD-1 therapy. **(A)** Heatmap showing normalized expression (log2 fold-change, mean-centered per gene) of a 15-gene IFN and innate immune pathway stimulation (IIPS) signature across cutaneous (CM), acral (AM), and mucosal (MM) melanoma PDX tumors from the CU cohort. **(B)** Average IIPS signature scores by melanoma subtype. **(C)** IIPS scores by anti-PD-1 and/or anti-CTLA-4 combined with anti-PD-1 therapy response, where response was defined as any evidence of disease control as determined by RECIST v1.1 criteria or by treating physician assessment. **(D)** CIBERSORT immune deconvolution analysis of PDX RNA-seq data. Models in the top 20% (IIPS-high) or bottom 20% (IIPS-low) based on IIPS signature scores were compared. \**P* < 0.05; \*\**P* < 0.01.

### Decitabine induces IIPS and adaptive immune gene expression in rare melanoma cell lines

To investigate strategies for restoring the IIPS pathway, we tested whether IIPS genes could be pharmacologically upregulated in IIPS low/deficient rare melanoma cell lines. We evaluated the expression of key IIPS pathway genes in human melanoma cell lines and confirmed lower IIPS gene expression in rare melanoma cells, consistent with the transcriptional profiles observed in tumor samples. Expression of *DDX58* (encoding RIG-I), and *IFI44* and *IFI6* (IFN response genes), was low in two mucosal and two uveal melanoma cell lines compared to two cutaneous melanoma lines. Among three acral melanoma lines, expression was variable, with one exhibiting high baseline levels and two showing low expression (Figure 4A).

**Figure 4.**
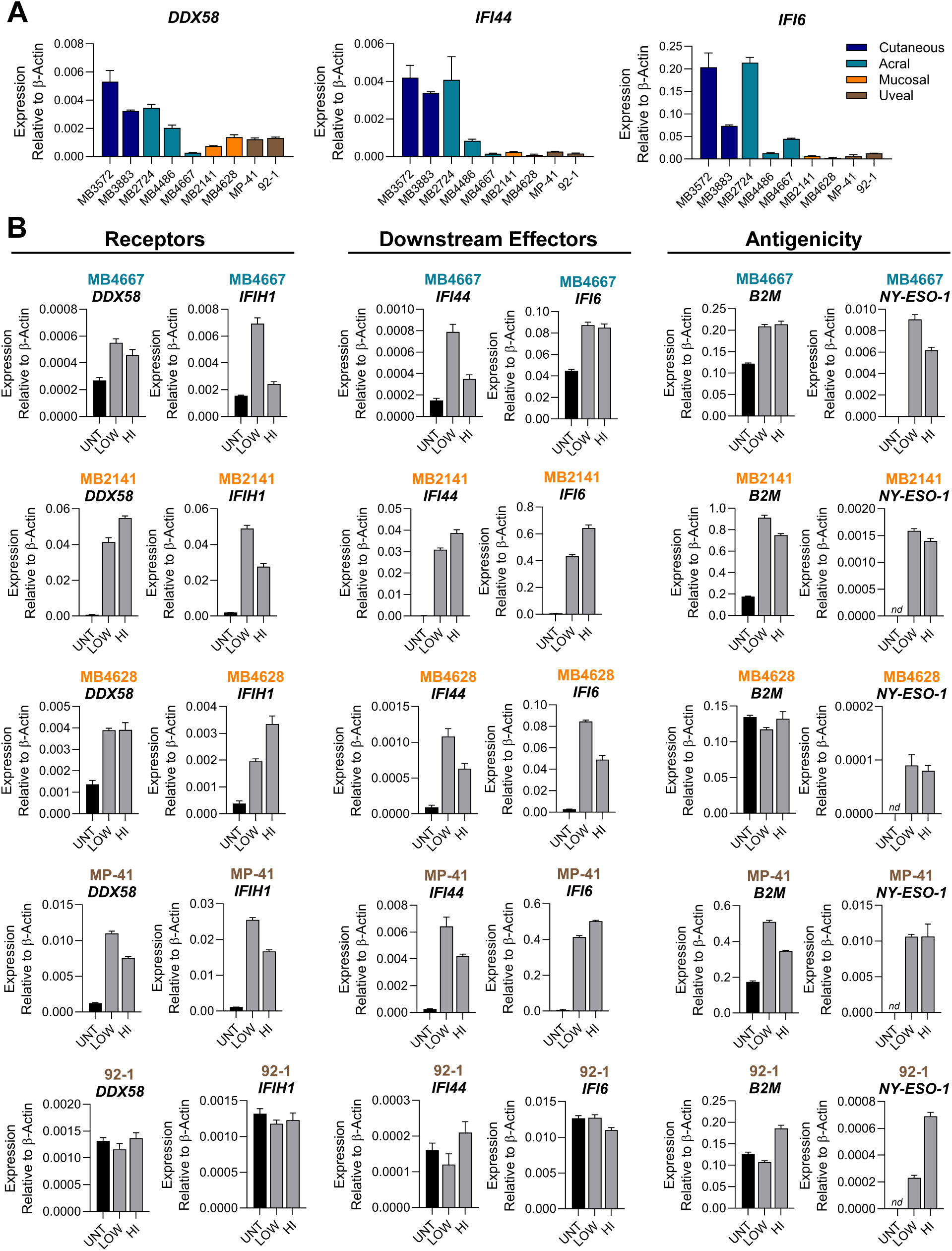
Decitabine induces IIPS and adaptive immune gene expression in rare melanoma cell lines. **(A)** Baseline expression of innate immune genes (*DDX58*, *IFI44*, *IFI6*) in melanoma cell lines representing cutaneous (blue), acral (teal), mucosal (orange), and uveal (brown) subtypes. Expression was measured by qRT-PCR and normalized to *β*-Actin. Error bars represent the standard error of the mean (SEM) from technical replicates. **(B)** Gene expression in rare melanoma cell lines following treatment with low-dose (LOW, 500 nM) or high-dose (HI, 1 µM) decitabine for four days. Cells were harvested 72 hours after the final treatment for qRT-PCR analysis. Expression levels of innate immune genes (*DDX58*, *IFIH1*, *IFI44*, *IFI6*) and adaptive immune genes (*B2M*, *NY-ESO-1*) were measured and normalized to *β*-Actin. UNT = untreated control. Error bars represent SEM of technical replicates. “nd” = not detected.

We tested four classes of drugs previously reported to activate the IIPS pathway in melanoma and other cancers (hypomethylating agents, CDK4/6 inhibitors, PARP inhibitors, and MYC inhibitors) using two agents from each class.^30–33^ These drugs were evaluated for their ability to induce *DDX58* and other IIPS response genes in two melanoma cell lines (Supplementary Figure 3). CDK4/6 and MYC inhibitors led to modest induction of all three genes at higher concentrations, while PARP inhibitors failed to induce expression of any IIPS genes (Supplementary Figures 3B-D). In contrast, hypomethylating agents robustly upregulated IIPS genes at both low and high doses, with decitabine outperforming 5-azacytidine, consistent with prior studies (Supplementary Figure 3A).^34^ In the cutaneous melanoma cell line, decitabine induced approximately threefold increases in IIPS gene expression. However, in the mucosal melanoma cell line, which had lower baseline expression, decitabine treatment led to more than 20-fold induction of all three genes (Supplementary Figure 3A).

Further testing confirmed that decitabine induced expression of IIPS genes (*DDX58, IFIH1, IFI44, IFI6*) and adaptive immune genes (*B2M, NY-ESO-1*) in four of five rare melanoma cell lines (Figure 4B). The only non-responder, the 92-1 uveal melanoma line, still demonstrated increased expression of adaptive immune genes despite minimal induction of IIPS genes (Figure 4B).

We then tested decitabine in xenograft models (MB2141 mucosal and MP-41 uveal) treated with prior tetrahydrouridine to enhance decitabine bioavailability (Figure 5A). Analysis of tumors showed that decitabine led to increased expression of IIPS and adaptive immune genes at 21 days post-treatment in the mucosal melanoma model and 14 days in the uveal melanoma model (Figure 5B). The timing of gene induction was consistent across all genes except for *IFI6,* which was induced as early as 7 days, and *MAGEA1* which continued to be increased out to 21 days, in the uveal melanoma model.

**Figure 5.**
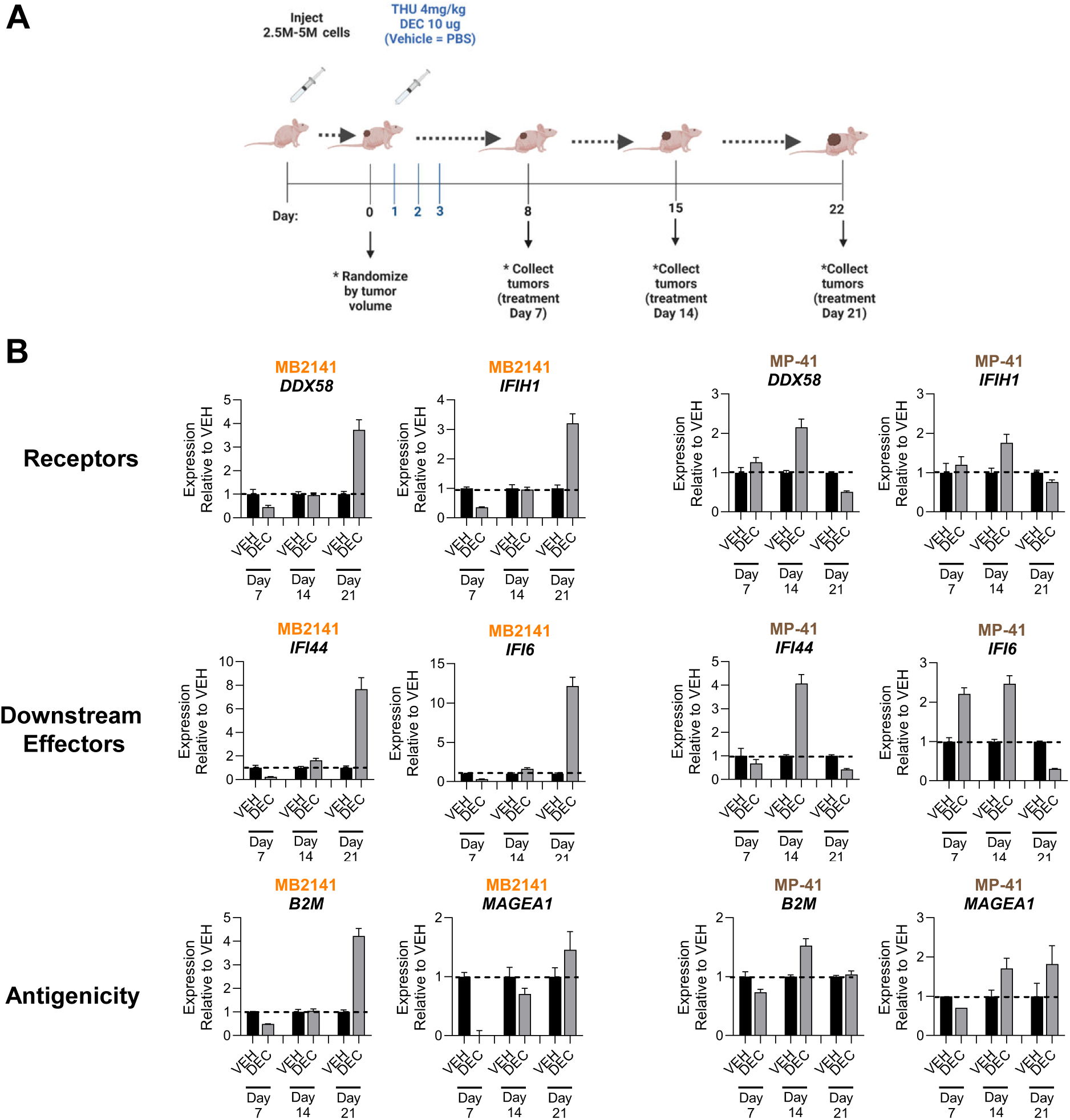
Decitabine induces innate and adaptive immune gene expression in rare melanoma xenograft models. **(A)** Schematic of *in vivo* treatment schedule. Mice bearing mucosal (MB2141) or uveal (MP-41) melanoma xenografts averaging 100-300 mm^3^ were randomized by tumor volume and treated subcutaneously once daily for three consecutive days with tetrahydrouridine (THU, 4 mg/kg) followed 30 minutes later by decitabine (DEC, 10 μg per mouse). Tumors were harvested at days 7, 14, or 21 post-treatments. **(B)** Expression of innate immune genes (*DDX58, IFIH1, IFI44, IFI6*) and adaptive immune genes (*B2M, MAGEA1*) in DEC-or vehicle (VEH)-treated tumors at each timepoint, measured by qRT-PCR. Expression values were normalized to *β*-Actin and plotted relative to VEH controls for each timepoint. Error bars represent the standard error of the mean (SEM) of technical replicates.

### Decitabine globally restores IIPS gene expression and activates IFN response programs

To better understand the broader transcriptional effects of decitabine and determine whether IIPS induction reflected a targeted or global shift in tumor gene expression, we performed whole-transcriptome sequencing of treated cells and tumors. RNA sequencing of decitabine-treated MB2141 mucosal melanoma cells and xenografts (Supplementary Data 3) revealed robust upregulation of IIPS and IFN response genes (Figure 6A), including the cytosolic sensors *DDX58* and *IFIH1*, as well as downstream effectors *IFI6*, *IFI27*, and *IFI44*. In parallel, we observed increased expression of antigen-presentation genes such as *B2M*, *TAP1*, and *HLA-A*, *HLA-B*, and *HLA-C*. Pathway enrichment analysis of upregulated transcripts (Supplementary Data 4) showed that decitabine treatment led to near-exclusive activation of IIPS and IFN-related pathways (Figure 6B). Thus, decitabine treatment selectively restores the expression of key IIPS and antigen-presentation genes and produces a broad IFN-driven transcriptional response.

**Figure 6.**
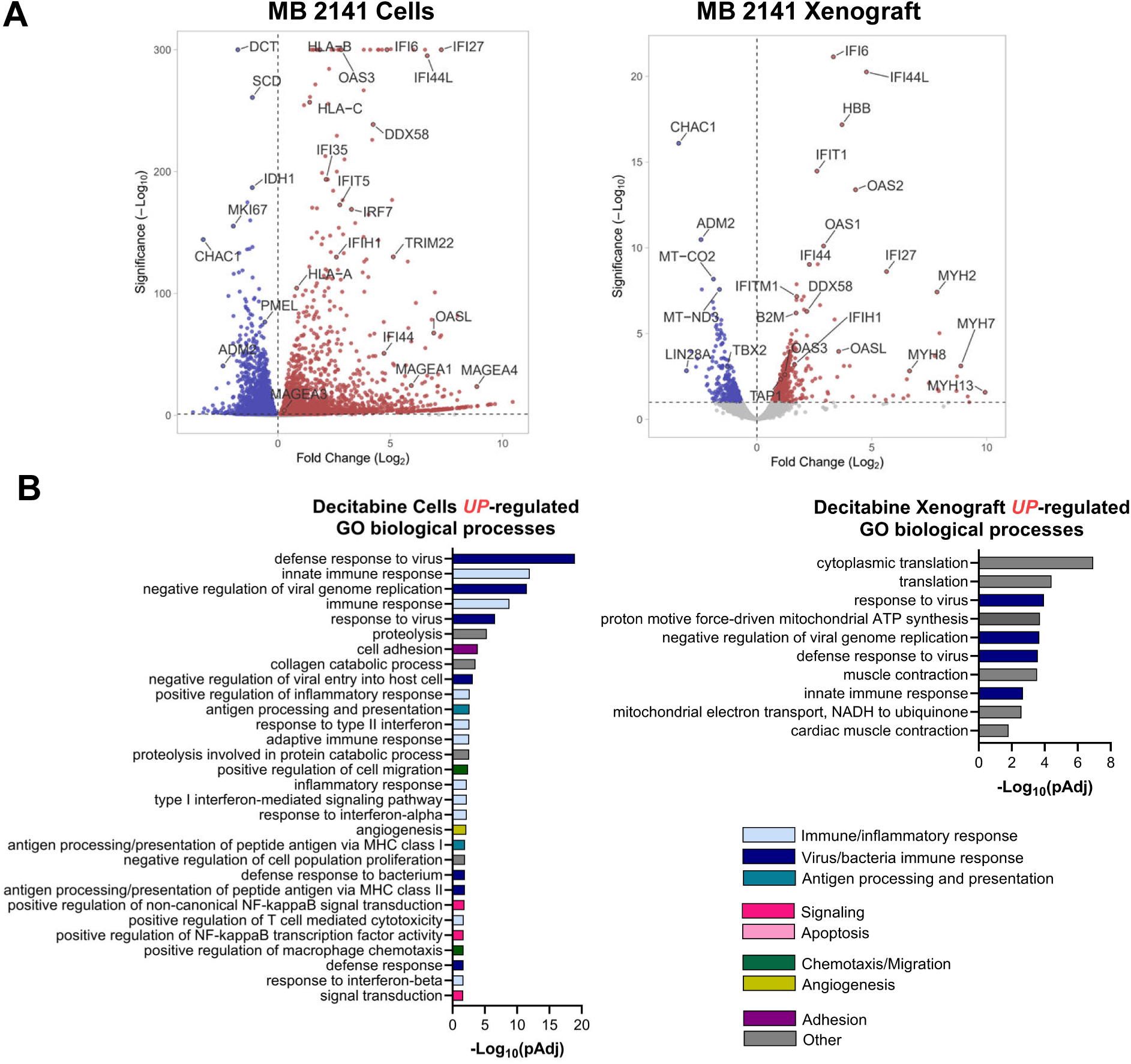
Decitabine selectively induces IIPS pathway genes in rare melanoma. **(A)** Volcano plots showing differentially expressed genes between untreated and decitabine-treated MB2141 cells (left) and xenografts (right). Genes with adjusted *P*-values < 0.05 and absolute log₂ fold change > 1 are highlighted. Key upregulated genes involved in IIPS and IFN signaling (*DDX58, IFIH1, IFI6, IFI44, IFI27, MAGEA1*) are labeled. **(B)** Top Gene Ontology (GO) biological processes significantly enriched (*P* < 0.05, FDR-adjusted) among upregulated genes in decitabine-treated MB2141 cells (left) and xenografts (right). Bars are color-coded by functional category, highlighting predominant enrichment of innate immune, IFN, and antigen processing pathways *in vitro*, with a more heterogeneous response observed *in vivo*.

## Discussion

The effectiveness of immune checkpoint inhibitors (ICIs) in melanoma has been primarily defined by studies of cutaneous melanoma, where response rates to anti-PD-1 and anti-CTLA-4 inhibitors are among the highest of any solid tumor.^1,2^ In contrast, rare melanoma subtypes (including acral, mucosal, and uveal) exhibit far poorer responses,^5–14^ with limited understanding of the molecular drivers of resistance. In this study, we performed a comparative transcriptomic analysis of cutaneous and rare melanomas and identified consistent downregulation of the innate immune pathogen-sensing (IIPS) pathway as a defining feature of rare melanomas, particularly mucosal and uveal melanomas. We further show that pharmacologic reactivation of this pathway with the hypomethylating agent decitabine restores IIPS gene expression and enhances tumor immunogenicity in preclinical models. These findings provide insight into a shared, targetable mechanism of immune resistance in rare melanomas.

The role of the IIPS pathway in shaping anti-tumor responses is an area of increasing interest in the field of immuno-oncology. Cytosolic pattern recognition receptors, such as RIG-I (*DDX58*) and MDA5 (*IFIH1*), detect viral-like RNA species within tumor cells and trigger downstream type I IFN signaling.^35^ This so-called “viral mimicry” response has been shown to play an important role in priming adaptive immunity and enhancing responsiveness to checkpoint blockade.^36,37^ In melanoma specifically, preclinical studies have demonstrated that tumor cell-intrinsic IIPS signaling is critical for response to ICI.^38,39^ Our transcriptomic analysis is the first to reveal significantly lower expression of IIPS-related genes in mucosal and uveal melanomas compared to cutaneous melanoma, and this suppression was also observed in acral melanoma, albeit to a lesser extent. Notably, tumors from anti-PD-1 non-responders in our PDX cohort also exhibited low IIPS gene expression. Immune deconvolution analysis further revealed that tumors with low IIPS expression were enriched for immunosuppressive cell populations, including M2 macrophages, and depleted of effector CD8⁺ and memory CD4⁺ T cells, suggesting that IIPS silencing may contribute to a non-inflamed tumor microenvironment. These findings support a model in which transcriptional silencing of the IIPS pathway may serve as a shared mechanism of immune evasion across rare melanoma subtypes.

We further demonstrate that low IIPS gene expression in rare melanomas can be restored pharmacologically *in vitro* and *in vivo* by treatment with decitabine, a known inducer of IIPS/type I IFN gene programs.^36,40–43^ Decitabine has traditionally been used to treat hematologic malignancies, including acute myeloid leukemia and chronic myelomonocytic leukemia,^44^ but recent studies have explored its immunomodulatory potential in lymphoma and solid tumors.^45,46^ Preclinical models of melanoma, colorectal cancer, and triple-negative breast cancer have shown that decitabine enhances responses to anti-PD-1 therapy.^47–50^ In early-phase clinical trials, combinations of hypomethylating agents with ICIs have shown promising activity. For example, guadecitabine plus pembrolizumab achieved a 37% disease control rate in a phase I study in patients with advanced solid tumors.^46^ In relapsed/refractory Hodgkin lymphoma, decitabine combined with camrelizumab resulted in a 71% complete response rate, more than double that of camrelizumab alone.^45^ These data have led to a growing number of clinical trials exploring hypomethylating agents in solid tumors, including our own clinical trial (NCT05089370) evaluating oral cedazuridine (decitabine/tetrahydrouridine) in combination with nivolumab in patients with unresectable stage III or IV mucosal melanoma.

This study has several limitations. We used high-quality RNA from PDX tumors to characterize acral and mucosal transcriptomes. While PDX models retain many key features of the originating tumors, including histopathology, gene expression, and epigenetic landscapes,^51^ they lack tumor-infiltrating immune cells and may not fully recapitulate the tumor microenvironment. However, the observation of IIPS downregulation was independently validated in the TCGA uveal melanoma cohort, which includes direct patient tumor samples not PDX tumors, lending clinical relevance to our findings. Another limitation is the relatively small number of rare melanoma samples with known ICI treatment responses, which may limit the generalizability of our immune correlation findings. Additionally, our study primarily evaluated the transcriptional consequences of decitabine treatment, and further studies are needed to define its effects on protein expression, immune cell recruitment, and functional antitumor immunity *in vivo*. Finally, while we used CIBERSORT to infer immune cell composition, these estimates rely on bulk RNA-seq data and may not capture spatial or functional heterogeneity within the tumor microenvironment.

In summary, we identify innate immune signaling, specifically the IIPS/type I IFN pathway, as a shared, tumor-intrinsic vulnerability across rare melanoma subtypes. Our data suggest that epigenetic reactivation of this pathway using decitabine may help restore tumor immunogenicity and overcome resistance to immunotherapy. These findings provide a mechanistic foundation for ongoing and future trials that combine hypomethylating agents with ICI in patients with acral, mucosal, and uveal melanoma.

## Research Support

Supported by CDMRP MRP Award W81XWH-21-1-0660, NIH NCI P30CA046934, the CU Center for Rare Melanomas funding by the Patten-Davis Foundation, and the Moore Family Foundation.

## Acknowledgements

We first and foremost thank the patients for consenting to use their clinical data and specimens for this study. We thank the National Jewish Health Genomics Core Facility for sequencing support. This work was supported by CDMRP MRP Award W81XWH-21-1-0660, NIH NCI P30CA046934 CU Cancer Center Support Grant, the CU Center for Rare Melanomas funding by the Patten-Davis Foundation, and the Moore Family Foundation.

## Author Contributions

K.L. Couts conceived and designed the study. M.L. MacBeth, S.M. Bagby, J.A. Turner, and P.A. Whitty collected data and performed experiments. K.C. Anderson conducted formal RNA-seq data analysis. J.S.W. Borgers, R.P. Tobin, and K.L. Couts wrote the original draft. The manuscript was reviewed and edited by V.W. Rebecca, E. Davila, T.M. Pitts, T.M. Medina, S.P. Patel, M.D. McCarter, W.A. Robinson, and K.L. Couts. W.A. Robinson, R.P. Tobin, and K.L. Couts supervised the project.

## Data Availability Statement

The PDX RNA-seq dataset is available through dbGaP (accession number phs002951.v1.p1). RNA-seq data from decitabine-treated cell lines and xenograft tumors are being deposited in the NCBI Gene Expression Omnibus (GEO) and will be made publicly available upon publication. All other data supporting the findings of this study are included in the manuscript and supplementary materials.

