## Supplementary Figures for "Decitabine reverses innate immune gene suppression in rare melanomas"

### Supplementary Figure 1

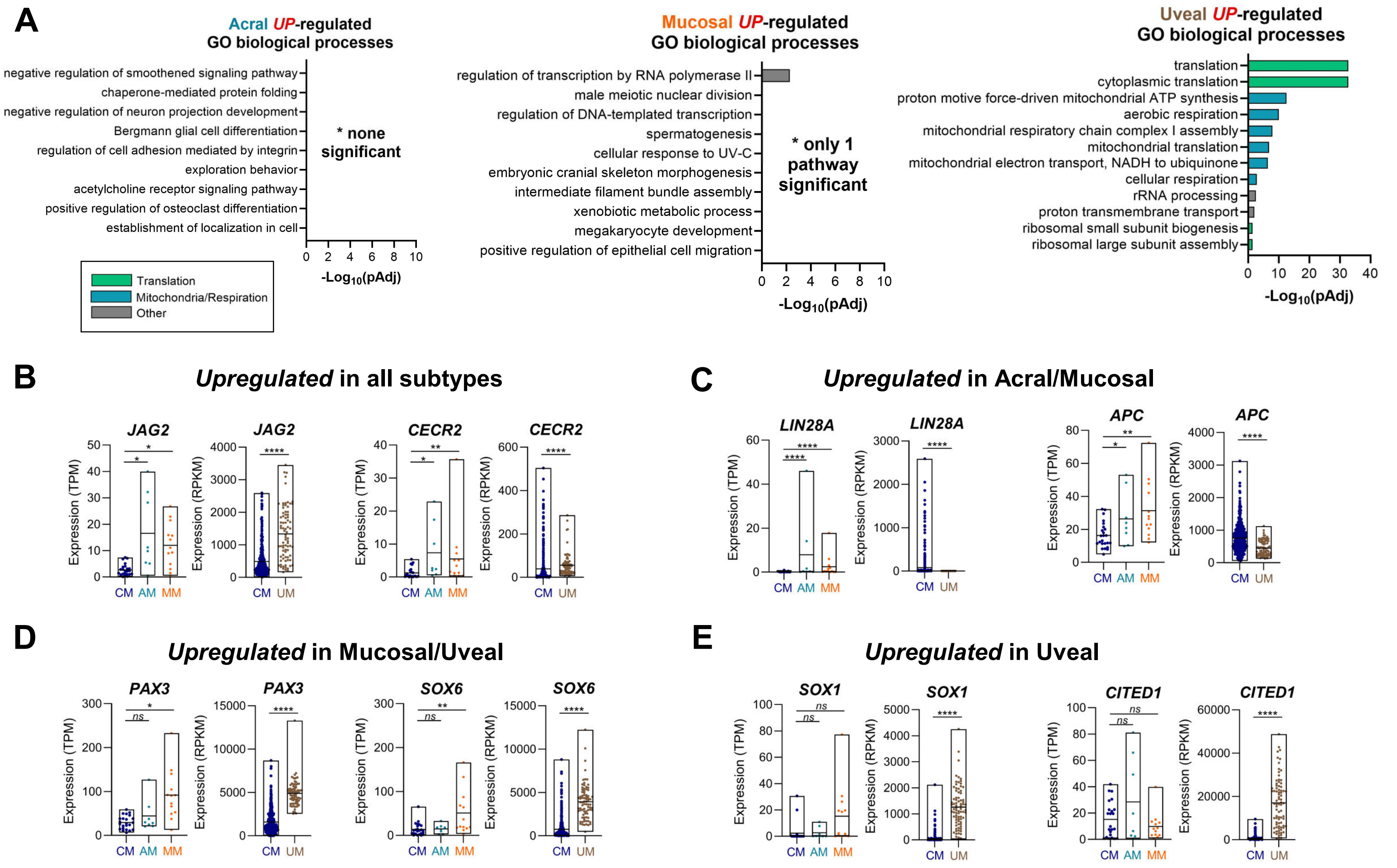


**Supplementary Figure 1. Up-regulated pathways and genes in distinct melanoma subtypes. (A)** Gene Ontology (GO) biological process enrichment analysis for genes significantly upregulated in acral (left), mucosal (middle), and uveal (right) melanomas compared to cutaneous melanomas. Bars are color-coded by category, including translation, mitochondrial/respiratory, and other biological processes. No pathways were significantly enriched for acral melanoma, and only one pathway was enriched for mucosal melanoma. **(B–E)** RNAseq expression of selected genes significantly upregulated in acral **(B)**, mucosal **(C)**, and uveal **(D, E)** melanoma subtypes, with corresponding values in cutaneous melanoma for comparison. Expression data are shown as TPM (PDX dataset) or RPKM (TCGA dataset), as indicated. Statistical significance is derived from the differential expression analysis shown in Supplementary Data 1. **** $p$<0.0001, *** $p$<0.001, ** $p$<0.01, $p$<0.05, ns = not significant.

#### Supplementary Figure 2

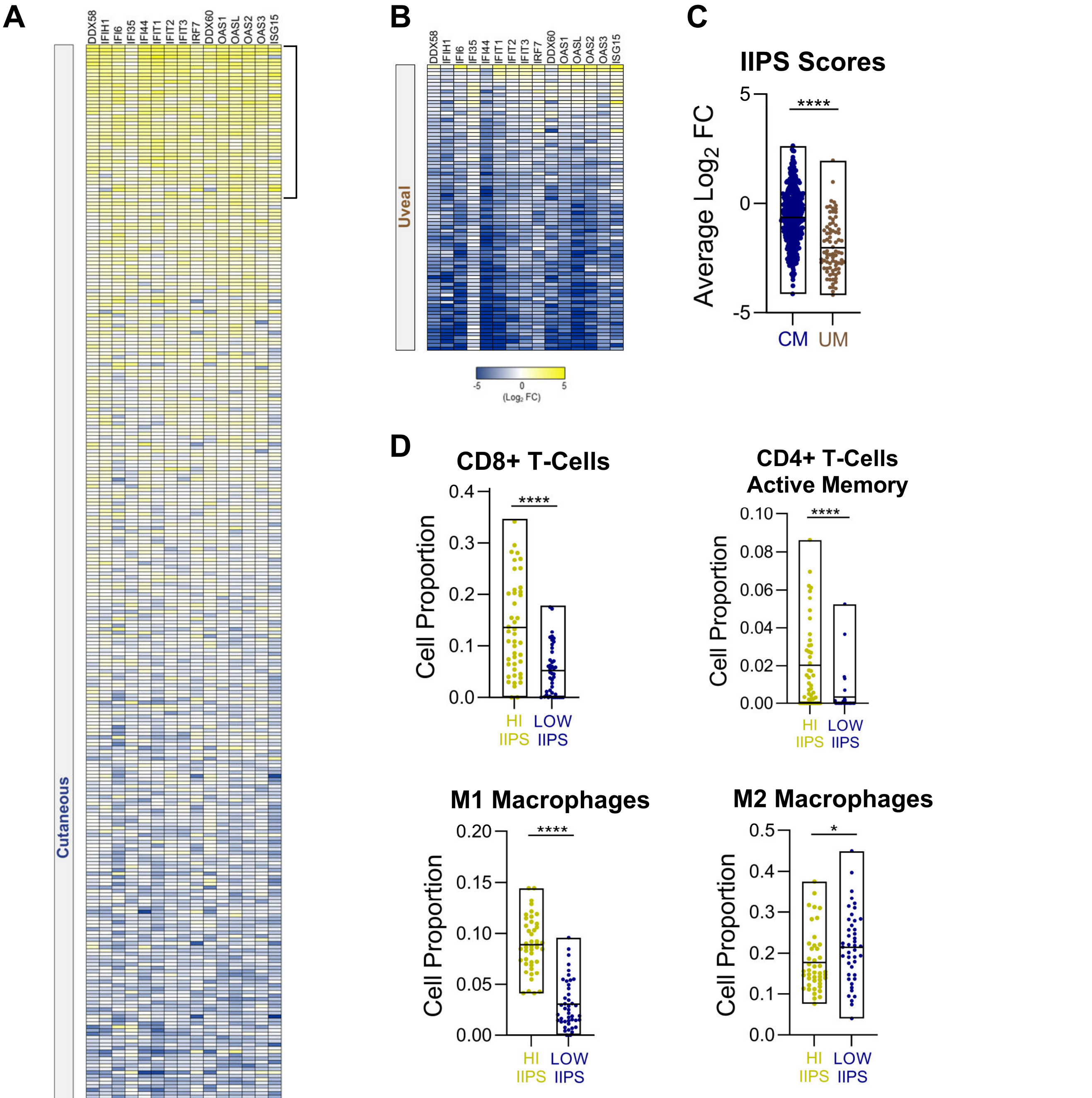


**Supplementary Figure 2. IIPS signature is low in UM and correlates with predicted immune infiltration in CM. (A-B)** Heatmap showing expression (mean-centered, log2-transformed) of 15 selected type I interferon and innate immune pathway stimulation (IIPS) genes in **(A)** cutaneous melanoma or **(B)** uveal melanoma tumors from The Cancer Genome Atlas (TCGA-SKCM). Genes were selected based on consistent upregulation in decitabine-treated samples and/or known function in IIPS. **(C)** Average IIPS signature scores (computed as the mean of log2 fold-changes for all 15 genes) **(D)** Immune cell population estimates from CIBERSORT analysis of TCGA cutaneous melanoma tumors. Tumors in the top 10% (IIPS-high) and bottom 10% (IIPS-low) based on IIPS signature score were compared. (*p < 0.05, ****p < 0.0001, unpaired t-test).

### Supplementary Figure 3

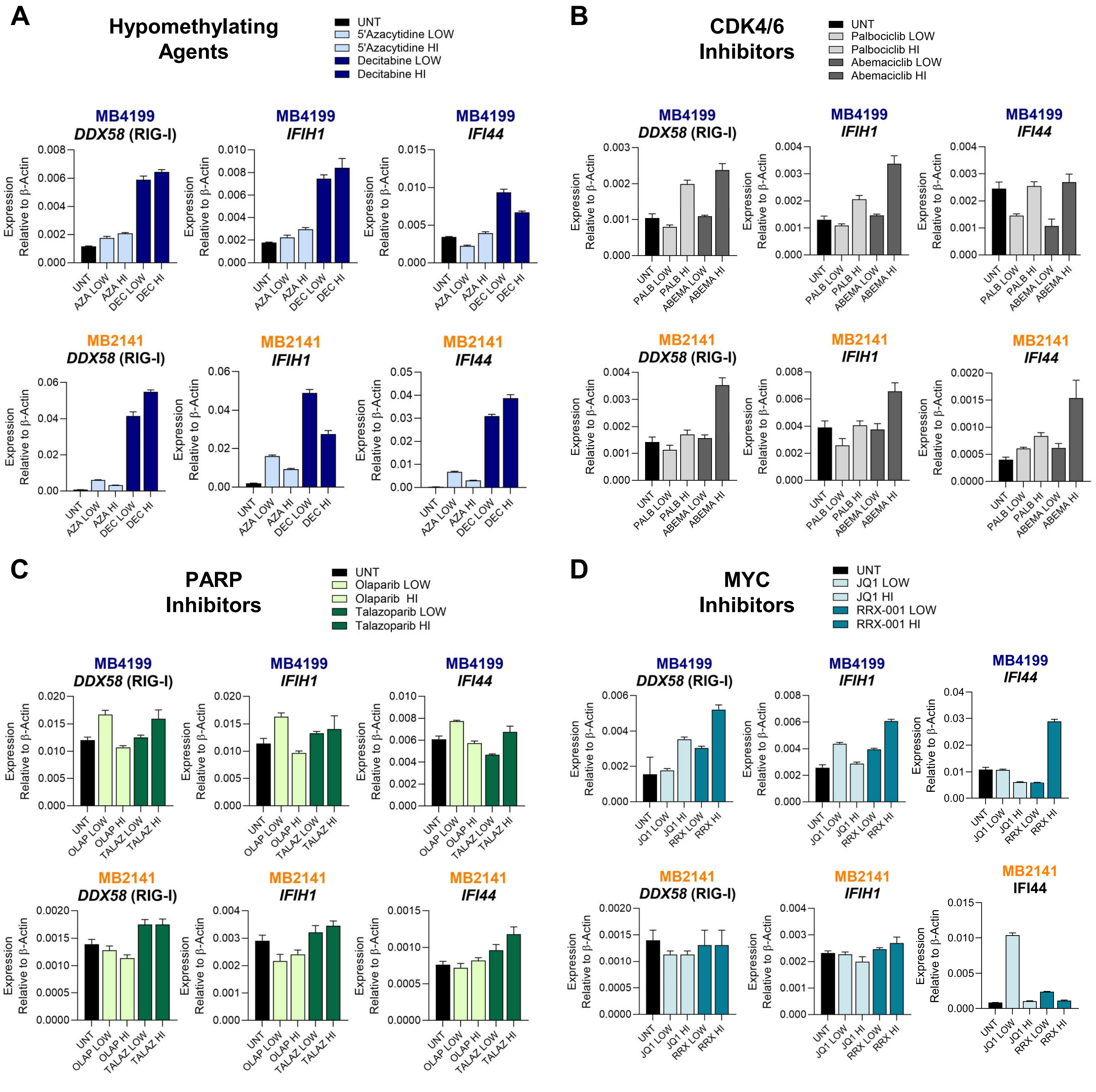


**Supplementary Figure 3. Drug screening identifies decitabine as a potent inducer of type I IFN and IIPS gene expression. (A–D)** Gene expression of *DDX58* (RIG-I), *IFIH1* (MDA5), and *IFI44* was evaluated by qRT-PCR in acral (MB4199) and mucosal (MB2141) melanoma cell lines following treatment with candidate compounds. Expression was normalized to β-Actin. Error bars represent the standard error of the mean (SEM) from technical replicates. **(A)** Cells were treated with low (500 nM) or high (1 μM) doses of 5-azacytidine (AZA) or decitabine (DEC) for 3 consecutive days followed by 4 days without treatment. **(B)** Cells were treated with low (500 nM) or high (1 μM) doses of palbociclib or abemaciclib for 48 hours. **(C)** Cells were treated with low (2.5 μM) or high (10 μM) doses of olaparib or talazoparib for 48 hours. **(D)** Cells were treated with low (100 nM) or high (500 nM) doses of JQ1, or low (500 nM) or high (2 μM) doses of RRX-001 for 48 hours.
