## Supplementary Methods for "Decitabine reverses innate immune gene suppression in rare melanomas"

*Whole transcriptome sequencing of PDX models*

cDNA libraries were prepared with the KAPA Stranded RNA-Seq Kit with RiboErase (KAPA Biosystems) per the manufacturer's protocol for Illumina systems. Input was 300 ng RNA, fragmentation was performed for 8 minutes at 94ºC to a size of 100-200 nucleotides, final adapter concentration of 15 nM, and 11 PCR cycles for amplification. Libraries were sequenced on a HiSeq2500 (Illumina), with five samples multiplexed per lane. Obtained sequencing reads were paired-end and 125 bp in length.

*Whole transcriptome data analysis of PDX models*

Raw reads were mapped to the human genome (hg38) with STAR aligner.[1] The Subread feature counts tool[2] was used to generate gene-level counts (Supplementary Data 5). Differential gene expression was called using DESeq2[3] using Wald tests. The p-values were corrected for multiple comparisons using the false-discovery rate method by Benjamini and Hochberg.[4] Volcano plots were generated using VolcaNoseR, and pathway enrichment analysis was performed using the DAVID functional annotation tool.[5,6] Heatmaps were generated using the Heatmapper web tool.[7]

*Analysis of TCGA data*

The gene-expression quantification tables (HTseq-count versions) for the TCGA cutaneous and uveal samples were downloaded using the Genomic Data Commons Application Programming Interface (API)[8]. The counts tables were all downloaded in March of 2019 (Supplementary Data 6). Differential gene expression was called using DESeq2[3] using Wald tests. The p-values were corrected for multiple comparisons using the false-discovery rate method by Benjamini and Hochberg.[4]

*RNA sequencing of decitabine-treated cells and xenograft tumors*

RNA sequencing libraries were prepared using the NEBNext Ultra II RNA Library Prep Kit for Illumina using the manufacturer's instructions (NEB, Ipswich, MA, USA). The sequencing libraries were clustered on a flow cell. After clustering, the flow cell was loaded on the Illumina NovaSeq instrument according to the manufacturer's instructions. The samples were sequenced using a 2x150bp Paired-End (PE) configuration, targeting 30 M reads/sample. Raw sequence data (.bcl files) generated by the sequencer were converted into FASTQ files and de-multiplexed using Illumina's bcl2fastq 2.20 software. Sequence reads were trimmed to remove possible adapter sequences and nucleotides with poor quality. The trimmed reads were mapped to the reference genome available on ENSEMBL using the STAR aligner v.2.5.2b[1]. Unique gene hit counts were calculated by using feature Counts from the Subread package v.1.5.2[2]. Using DESeq2[3], a comparison of gene expression between the groups of samples was performed (Supplementary Data 3). The Wald test was used to generate *p*-values and Log2 fold changes.

*Quantitative Real-Time PCR Primer sequences*

| Primer | Sequence, 5' → 3' |
|---|---|
| β-Actin Forward<br>β-Actin Reverse | AGAGCTACGAGCTGCCTGAC<br>AGCACTGTGTTGGCGTACAG |
| DDX58 Forward<br>DDX58 Reverse | AAACCAGAATTATCCCAACCG<br>TGATCTGAGAAGGCATTCCAC |
| IFI44 Forward<br>IFI44 Reverse | TGGTACATGTGGCTTTGCTC<br>CCACCGAGATGTCAGAAAGAG |
| IFI6 Forward<br>IFI6 Reverse | CGCTTTTCTTGTGCTACCTG<br>TGTCCGAGCTCTCCGAG |

| | |
|---|---|
| IFIH1 Forward<br>IFIH1 Reverse | TTGATGGTCCTCAAGTGGAAG<br>CTTCTGCCAAACTTGTGTCTG |
| B2M Forward<br>B2M Reverse | AGGCTATCCAGCGTACTCCA<br>TCATCCAATCCAAATGCGGC |
| NY-ESO-1 Forward<br>NY-ESO-1 Reverse | TGCTTGAGTTCTACCTCGCCA<br>TATGTTGCCGGACACAGTGAA |
| MAGEA1 Forward<br>MAGEA1 Reverse | GCCAAGCACCTCTTGTATCCTG<br>GGAGCAGAAAACCAACCAAATC |

**References- Supplementary Methods**
